# *stably*: error-controlled stability selection for biomarker panel discovery

**DOI:** 10.64898/2026.09.11.750833

**Authors:** Daniel Byrne, Mairéad G McNamara, Richard D Unwin

**Affiliations:** Division of Cancer Sciences, School of Medical Sciences, Faculty of Biology, Medicine, and Health, The University of Manchester, Manchester, M13 9NY, United Kingdom; The Christie NHS Foundation Trust, Manchester M20 4BX, UK

**Keywords:** data-independent acquisition, mass spectrometry, clinical proteomics, biomarker discovery, stability selection, false discovery control, machine learning, pancreatic cancer

## Abstract

**Motivation:** Data-independent acquisition mass spectrometry (DIA-MS) has become increasingly popular for clinical proteomics due to its sensitivity and reproducibility. Univariate statistical analysis tools, such as *limma* and *MSstats*, are widely used for identifying differentially abundant proteins but cannot capture multivariate relationships between proteins that may provide greater discriminatory power as a panel. Machine learning approaches can address this gap, but typically prioritise predictive performance over feature stability, producing biomarker panels for downstream validation that vary depending on data splitting and are poorly suited to clinical translation.

**Results:** We developed *stably*, a Python package that implements stability selection with formal false positive control for DIA proteomics data. Using synthetic data with known ground-truth biomarkers, we show that the Shah and Samworth complementary pairs stability selection framework recovers more true synthetic biomarkers than the Meinshausen and Bühlmann framework at moderate effect sizes typical of proteomics (d = 0.5 – 2.0), while both maintain false positive rates well below their theoretical guarantees. Applied to a publicly available serum proteomics dataset from patients with all stages of pancreatic ductal adenocarcinoma (n=176), *stably* identified a stable 17-protein biomarker panel in the discovery cohort (n=120), which achieved higher predictive power (AUC = 0.93) in the validation cohort (n=56) than the panel selected by Byeon et al. (2024)(AUC 0.82). *stably* represents a principled, error-controlled method for biomarker panel discovery for translation into second cohorts.

**Availability and implementation:** *stably* is available on GitHub (https://github.com/byrnedaniel5-eng/stably); the version used in this study is archived on PyPI (https://pypi.org/project/stably/).

## Introduction

Biomarker discovery requires us to discern markers with high sensitivity and reproducibility to enable the development of assays for the detection of both common and rarer diseases on a population level, where imperfect tests can cause significant harm to patients. Data-independent acquisition (DIA) mass spectrometry (MS) has become an increasingly popular method for protein biomarker discovery (Krasny and Huang, 2020). Analysis packages such as *limma* and *MSstats* perform univariate statistical tests to identify proteins that are differentially abundant within a dataset (Choi, et al., 2014; Ritchie, et al., 2015). Although useful for univariate analysis, these approaches are unable to capture relationships between proteins that may provide greater discriminatory power when considered as a panel; a gap that multivariable machine learning (ML) methods are well-positioned to address.

Standard ML for biomarker discovery involves splitting data into training, validation, and test sets (Adams and Bittremieux, 2025). The model classifies experimental groups in the training set, the hyperparameters of the base learner are optimised on the validation set, and the performance evaluation is produced on the held-out test set. While robust for datasets with high numbers of samples, this approach is poorly suited to typical clinical proteomics discovery experiments. A cohort size of 100 patients with a standard split of 50/30/20 yields only 20 samples for a held-out test set. Performance metrics are thus highly sensitive to individual patients, producing potentially misleading estimates of panel performance. Acquiring cohorts sufficiently large to avoid these problems is both logistically and financially challenging, especially for less common diseases. A common workaround is therefore to use repeated cross-validation (CV) or nested CV, which produces more stable performance estimates from small datasets (Desaire, 2022). However, CV typically optimises for predictive accuracy, not feature consistency (how often a feature is selected across folds); the set of proteins selected may differ substantially between subsamples, even when overall classification performance remains similar. This is problematic for biomarker panel discovery, where the goal is not accurate prediction in the experimental cohort, but identification of a reproducible set of proteins that can be carried forward into independent cohorts.

To encourage feature stability, Meinshausen and Bühlmann introduced a model of stability selection using repeated random subsampling of a dataset (Meinshausen and Bühlmann, 2010). In this model, the user sets their threshold of stability, defined as the minimum selection probability for a feature across subsamples; recommended to be between 60 and 90%. Two assumptions are required for this error bound to hold: that the selection probabilities for all noise proteins are the same, and the signal proteins should have a higher selection probability than noise proteins. This approach was improved by Shah and Samworth (2013), using complementary pairs stability selection (CPSS) instead of random subsampling, which ensures that all samples are equally sampled. Importantly, CPSS requires neither of the assumptions in the Meinshausen-Bühlmann model to be satisfied.

Both Meinshausen–Bühlmann and Shah–Samworth used an L1-penalised (Lasso) logistic regression as the primary base learner. However, when features are highly correlated, as is common in proteomics where proteins may co-vary, Lasso tends to arbitrarily select one feature from a correlated group while discarding the rest. ElasticNet combines Lasso and L2 (Ridge) penalisation, retaining the ability to drive coefficients to zero for feature selection, while the L2 component encourages correlated features to be shrunk together rather than arbitrarily discarded (Zou and Hastie, 2005). This makes it a more suitable base learner for proteomics data.

Several stability selection implementations exist, including the *stabs* package in R, which supports both the Meinshausen–Bühlmann and Shah–Samworth models, the stability-selection package from scikit-learn (now deprecated), and the *sharp* R package which integrates CPSS with ElasticNet (Bodinier, et al., 2025; Hofner, et al., 2015). Using these packages for clinical proteomic analysis requires advanced coding expertise to perform application-specific data processing, such as filtering for proteotypic peptides and scaling of data, before feature selection by stability selection. To address this, we developed *stably*, the first Python-based package that directly performs error-controlled ML-driven feature selection on proteomic datasets to identify candidate biomarker panels. We benchmark *stably* using synthetic data with known ‘biomarkers’ across a range of effect sizes, comparing both the Meinshausen–Bühlmann and Shah–Samworth models for suitability in untargeted proteomics data. We additionally apply *stably* to a publicly available dataset from patients with pancreatic cancer, identifying stable predictors of disease that are highly predictive across multiple cohorts, and perform better than candidates selected by other analyses.

## Methods

### stably

The *stably* workflow is shown in Figure 1. *stably* accepts the report.pg_matrix output of the common DIA analysis tool DIA-NN, and all parameters are user-configurable. Proteins detected with fewer than two proteotypic peptides, samples with a high proportion of missing proteins, and proteins missing in more than a user-defined percentage of samples are removed sequentially. Once the candidate set is fixed, the data are subsampled and imputation, scaling, and feature selection are performed within each subsample to prevent further data leakage between subsamples.

**Figure 1.**
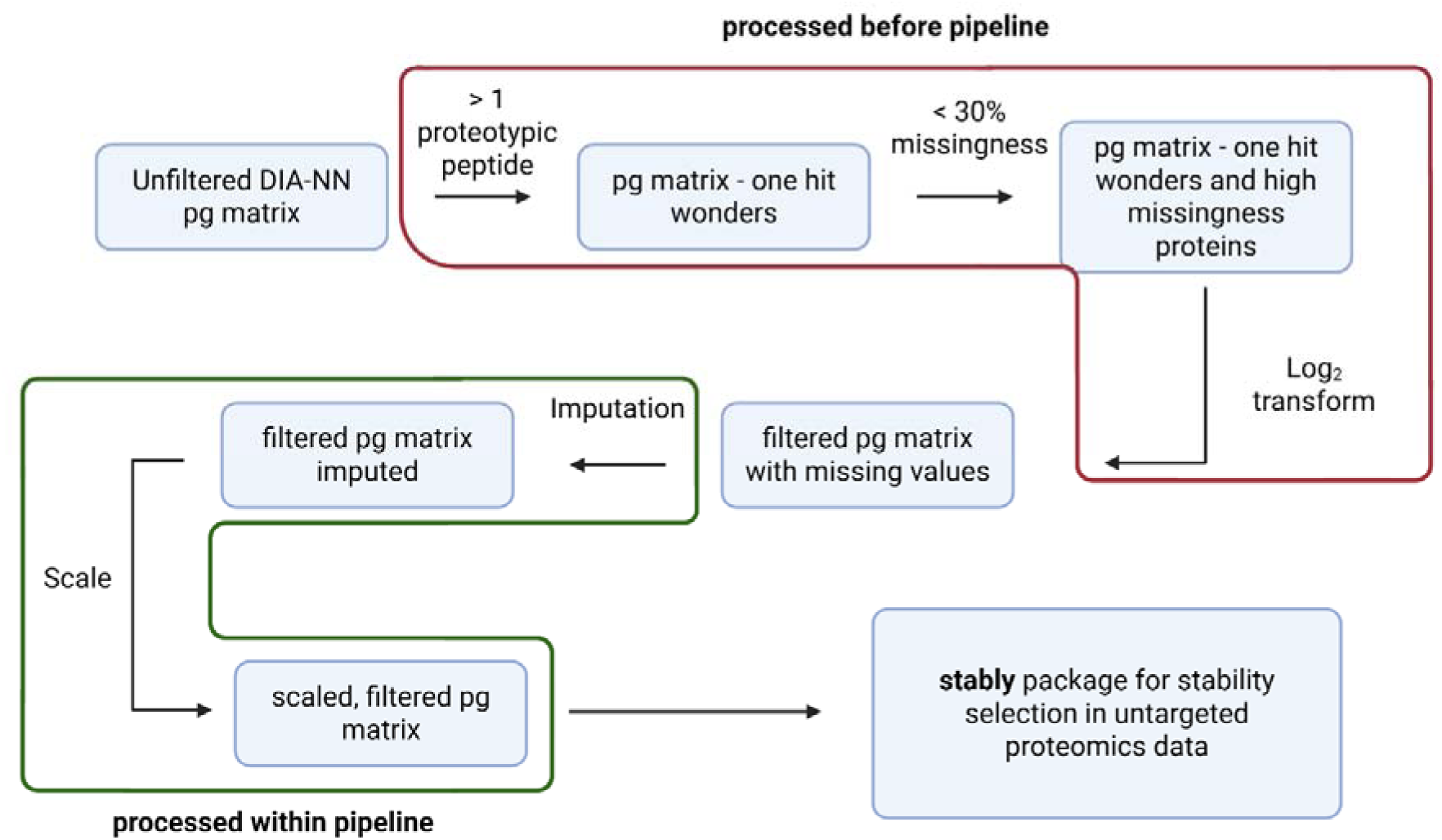
Flow diagram of the data-processing pipeline used by *stably* for biomarker discovery. The raw report.pg_matrix file from DIA-NN is read directly by *stably* and filtered before subsampling: proteins identified by fewer than two proteotypic peptides are removed, samples are filtered on protein coverage, and proteins with missingness above a user-defined threshold (default 30%) are then removed. This filtering precedes the split so that a fixed candidate set is present in every subsample — a requirement of both the Meinshausen–Bühlmann and Shah– Samworth false-positive bounds. The data are then subsampled, and imputation and standardisation are performed within each subsample to prevent leakage across subsamples. This ensures that the features selected in each subsample reflect only that subsample’s data, so that features passing the stability threshold are robustly stable candidates for downstream validation.

### Synthetic data generation

Synthetic datasets containing known differentially abundant proteins, were generated as previously described (Meyer, 2026). Baseline log_2_ protein intensities were sampled from a normal distribution (mean = 23, SD = 2) to match ranges observed in DIA proteomics data, with per-protein noise SDs drawn uniformly from 0.3–0.8 in log_2_ space. To display the ability of the synthetic data generator to produce realistic data, we generated synthetic data with identical numbers of samples and proteins to a dataset we later use to identify biomarkers of pancreatic cancer, specifically the discovery dataset outlined below (Byeon, et al., 2024). For the benchmarking of *stably* with synthetic data, each dataset contained 100 samples, with 50 samples in each group and 1000 proteins; 20 of which were randomly designated as differentially abundant with fold changes to achieve a target Cohen’s d. The Cohen’s d was fixed, with the required fold change back-calculated based on the noise within the dataset. Missing values were introduced whereby lower-abundance proteins exhibited higher missingness, mimicking the missing-not-at-random pattern observed in mass spectrometry data.

### Benchmarking of *stably*

For each target effect size (d = 0, 0.25, 0.5, 0.75, 1.0, 1.25, 1.5, 2.0, 2.5, and 3.0), independent datasets were generated, with the seeds held constant across effect sizes. With 10 repeats at each effect size, this resulted in 100 unique datasets, with the d = 0 runs acting as negative controls. A true positive (TP) was a designated biomarker retained as a stable feature, and a false positive (FP) as a non-designated biomarker retained as stable. Precision (TP/(TP+FP)), recall (TP/(TP+FN)), and F1 were computed for each dataset and averaged across the ten datasets per effect size (mean ± standard error). All benchmarking used an ElasticNet base learner with an L1 ratio of 0.5, weighting the L1 and L2 penalties equally, and a per-family error rate (PFER) target of 10. The Shah-Samworth and Meinshausen-Bühlmann models were compared on the same datasets, with a two-sided paired Wilcoxon signed-rank test at each effect size. Holm-corrected p values were reported for each effect size, as well as a single pooled paired test over a proteomics-typical effect size range (0.5 – 2.0).

### Stable biomarker discovery in pancreatic cancer cohorts to identify cross-cohort signals

The pancreatic cancer dataset was obtained from ProteomeXchange (PXD046438). This data contains a discovery cohort of 120 patients, of which 60 patients were diagnosed with pancreatic cancer, and a validation cohort of 56 patients, of which 28 patients were diagnosed with pancreatic cancer. More details on the clinicopathological characteristics of this cohort can be found in the original paper (Byeon, et al., 2024). The raw files for these cohorts were searched separately in DIA-NN (version 2.5.0) against a SwissProt human proteome (July 2025, 20,652 protein entries). The resulting report.pg_matrix was processed by *stably* to identify stable biomarkers of pancreatic cancer. Here, we used the Shah–Samworth model with a PFER target of 10, an ElasticNet base learner, and an L1 ratio of 0.5.

### Univariate differential abundance analysis

Univariate differential abundance analysis was performed independently for each cohort using *limma*. Protein groups identified by fewer than two proteotypic peptides and those missing in more than 30% of samples were removed. Intensities were log_2_(x + 1) transformed and missing values imputed by k-nearest neighbours (k = 5). A design matrix without an intercept was fitted by least squares and a disease-versus-control contrast assessed by empirical Bayes moderated t-test. Storey q-values were computed using the *qvalue* package, and proteins with q < 0.05 were considered significant. Concordance of protein rankings between cohorts was assessed using Spearman rank correlation of q-values (significance ranking) and of log_2_ fold changes (effect-size ranking).

### Composition of training and test data

Model training and feature selection was performed within the discovery cohort (120 patients: 60 PDAC, 60 controls; 214 protein groups after filtering). The Sydney validation cohort (56 patients: 28 PDAC, 28 controls) was reserved as an independent test set and was not used for feature selection, hyperparameter choice, imputation, scaling, or model fitting. *Stably* used complementary pairs stability selection with B = 50 complementary pairs (100 subsamples at 50% of the discovery cohort), an ElasticNet base learner with an L1 ratio of 0.5, at most q = 50 candidate features per subsample, and a PFER target of 10, giving a selection-probability threshold of 0.65 on the discovery cohort.

### Predictive validation of biomarker panels

Three panels were evaluated: the 17 proteins selected as stable in the discovery cohort, the two proteins selected as stable in both cohorts (GSN and APOA4), and the two-protein panel reported by Byeon et al. (2024) (PIGR and VWF). Models were trained on the discovery cohort only, using identical procedures. Protein intensities were log_2_-transformed and missing values filled with the median of that protein. Features were standardised to zero mean and unit variance, and an L2-penalised logistic regression classifier (scikit-learn, maximum 20,000 iterations) was fitted with pancreatic cancer as the positive class. The regularisation strength C was tuned separately for each panel by grid search over C = 10⁻□ to 10², maximising mean AUC under five-fold stratified CV repeated five times, with imputation and standardisation refitted within each training fold so that no information crossed folds during tuning. The selected values were C = 0.01 for the 17-protein panel and C = 0.1 for both two-protein panels.

The imputation medians, standardisation parameters and regression coefficients estimated on the discovery cohort were applied unchanged to the 56 validation samples, and the predicted probability of PDAC was used as the classifier score. Receiver operating characteristic (ROC) curves and areas under the curve (AUC) were computed against the true validation labels. ROC AUCs were compared using a paired bootstrap over the validation cohort: in each of 10,000 replicates, the 56 validation samples were resampled with replacement and the same resampled indices applied to both panels’ predicted probabilities, with the difference in AUC recorded, and replicates containing only one class discarded. Confidence intervals are the 2.5th–97.5th percentiles of the resulting distribution, and p-values were obtained by inversion. Model coefficients were held fixed during resampling, so the intervals reflect sampling variability in the validation cohort rather than variability in model training. A fixed random seed (42) was used throughout.

### Permutation testing

The significance of each stable panel was assessed by a label-permutation null. The label-independent stage of preprocessing – the proteotypic peptide filter, the missingness filter, and the log_2_ transformation – was fitted once on the full cohort and reused for every permutation. This ensured the number of candidate features and selection probability threshold remained fixed across each permutation. Within each permutation, the group labels were shuffled and the entire selection procedure repeated, including threshold calibration, complementary-pairs subsampling, and per-subsample imputation and standardisation. One thousand permutations were run per cohort, and the number of features reaching the stability threshold, as well as the largest selection probability, was recorded for each permutation. One-sided p values were computed as (1 + #{null ≥ observed}) / (1 + N), where N is the number of permutations, ensuring p values can never reach zero (Phipson and Smyth, 2010). Ninety-five percent confidence intervals are Clopper–Pearson intervals on the exceedance count.

## Results

### A synthetic benchmark captures realistic proteomic data properties while providing a conservative test of feature selection

Synthetic proteomic datasets were generated containing a known set of differentially abundant proteins planted at controlled effect sizes. To ensure relevance, key features of the synthetic data were compared to the publicly available serum proteomics dataset used for final proof-of-concept (PXD046438) containing 214 proteins across 120 samples after filtering; the synthetic data were generated at matched dimensions for comparison (Figure 2). The synthetic protein intensity distribution was unimodal with comparable, though narrower, dynamic range to the real data (Figure 2A). Missingness was intensity-dependent in both datasets (Figure 2B). The correlation network is dense in the real dataset, with 60.7% of proteins in the discovery cohort dataset strongly correlated (pairwise Pearson’s |r| > 0.5) with at least one other protein, compared with 1.3% in the synthetic dataset (Figure 2C). The synthetic data had higher dimensionality, with 27 principal components required to explain 50% of variance, compared to 11 in the real cohort (Figure 2D).

**Figure 2.**
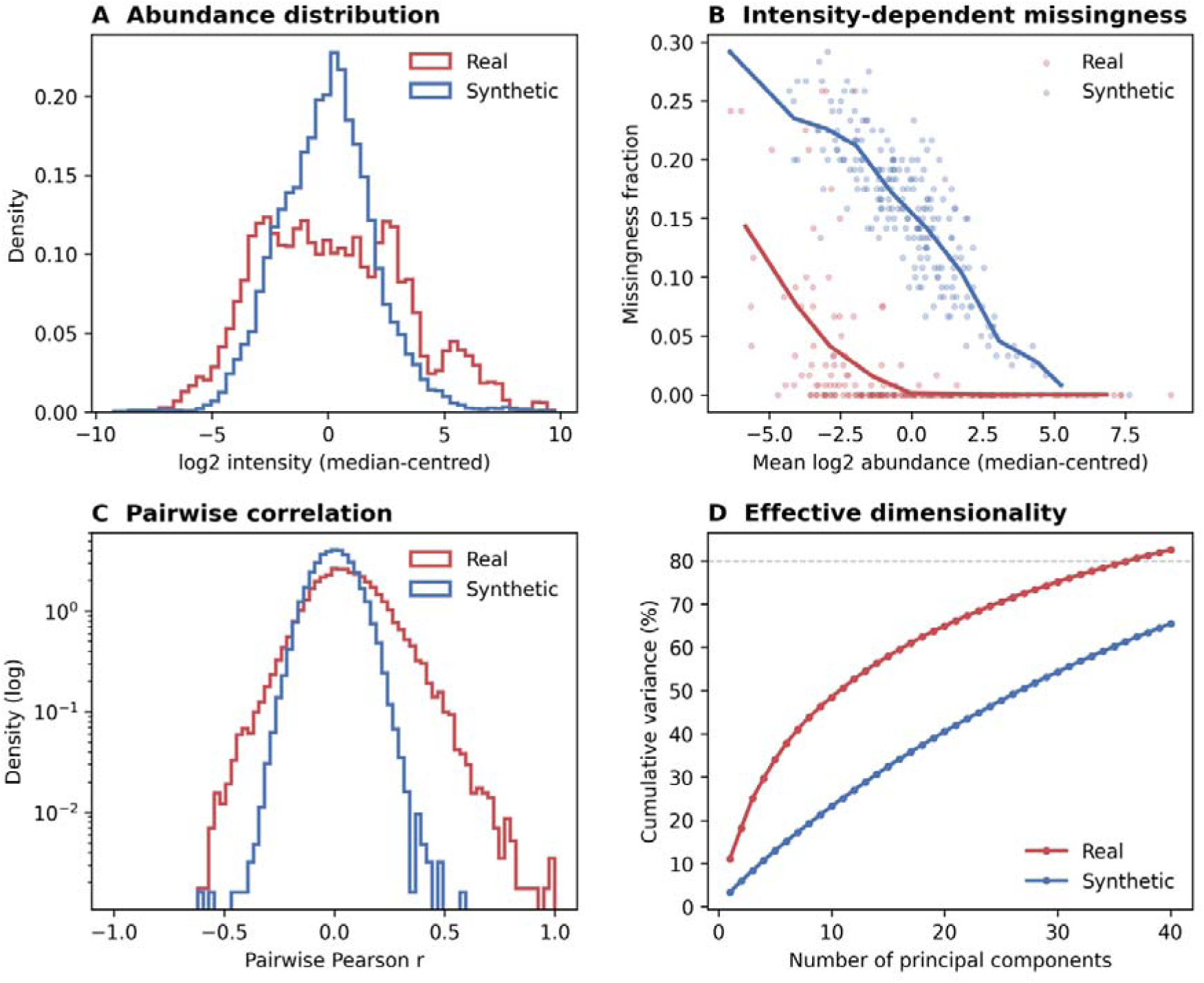
Characterisation of the synthetic benchmark against real serum proteomics. Real data: 214 proteins × 120 samples after filtering; synthetic data generated at matched dimensions. Absolute LFQ intensities are arbitrary, so distributions are median-centred. (A) log_2_ protein intensity distributions; the synthetic data reproduce a unimodal abundance distribution with a slightly narrower dynamic range. (B) Per-protein missingness versus mean abundance (points; lines = binned means): both datasets share the intensity-dependent (MNAR) missingness pattern. (C) Distribution of pairwise Pearson correlations (KNN-imputed, log-scaled density): real data have a substantially greater correlation network (D) Cumulative variance explained by each feature displays that the features within the synthetic data have less total variance loaded per protein than true data.

### Shah–Samworth theorem outperforms Meinshausen–Bühlmann on synthetic untargeted proteomics data at moderate effect sizes

To compare the performance of the Shah–Samworth and Meinshausen–Bühlmann models, we benchmarked both on synthetic untargeted proteomics data across a range of effect sizes using an ElasticNet base learner (L1 ratio = 0.5) at a PFER target of 10. At this target, the Shah–Samworth r-concave bound produced a lower selection-probability threshold than the Meinshausen–Bühlmann model (0.54 versus 0.71). This lower threshold translated into higher recall of true biomarkers at moderate effect sizes (Figure 3A). At d = 0.75, the Shah–Samworth model recovered 52% of true biomarkers versus 29% for Meinshausen–Bühlmann, and at d = 1.0, 85% versus 67% (Holm-adjusted paired Wilcoxon signed-rank, p < 0.05 at both). The two converged at higher effect sizes (95% versus 92% at d = 1.25, and approximately 98% for both from d = 1.5). Across the proteomics-typical range (d 0.5–2.0), the Shah–Samworth threshold recovered more true biomarkers than Meinshausen–Bühlmann (median recall difference +0.05; p = 8×10⁻□) and was never lower. This higher recall came at a modest cost to precision (Figure 3B): Meinshausen–Bühlmann achieved near-perfect precision (≥0.98 for d ≥ 0.75) by selecting fewer features, whereas Shah–Samworth precision ranged from 0.89 to 0.96 over the same range. The net effect on F1 favoured Shah–Samworth at moderate effect sizes (Figure 3C): F1 was significantly higher from d = 0.5 to 1.0 (0.65 versus 0.43 at d = 0.75; 0.89 versus 0.79 at d = 1.0; Holm-adjusted p < 0.05), converging at d ≈ 1.25 and crossing over at higher effect sizes, where Meinshausen– Bühlmann was marginally but significantly higher (0.95 versus 0.98 at d = 3.0; p < 0.05 at d ≥ 2.5). Both thresholds produced false positives far below their guaranteed bound (Figure 3D): mean false positives over d 0.5–2.0 were 1.0 for Shah– Samworth and 0.3 for Meinshausen–Bühlmann, against a PFER bound of 10.

**Figure 3.**
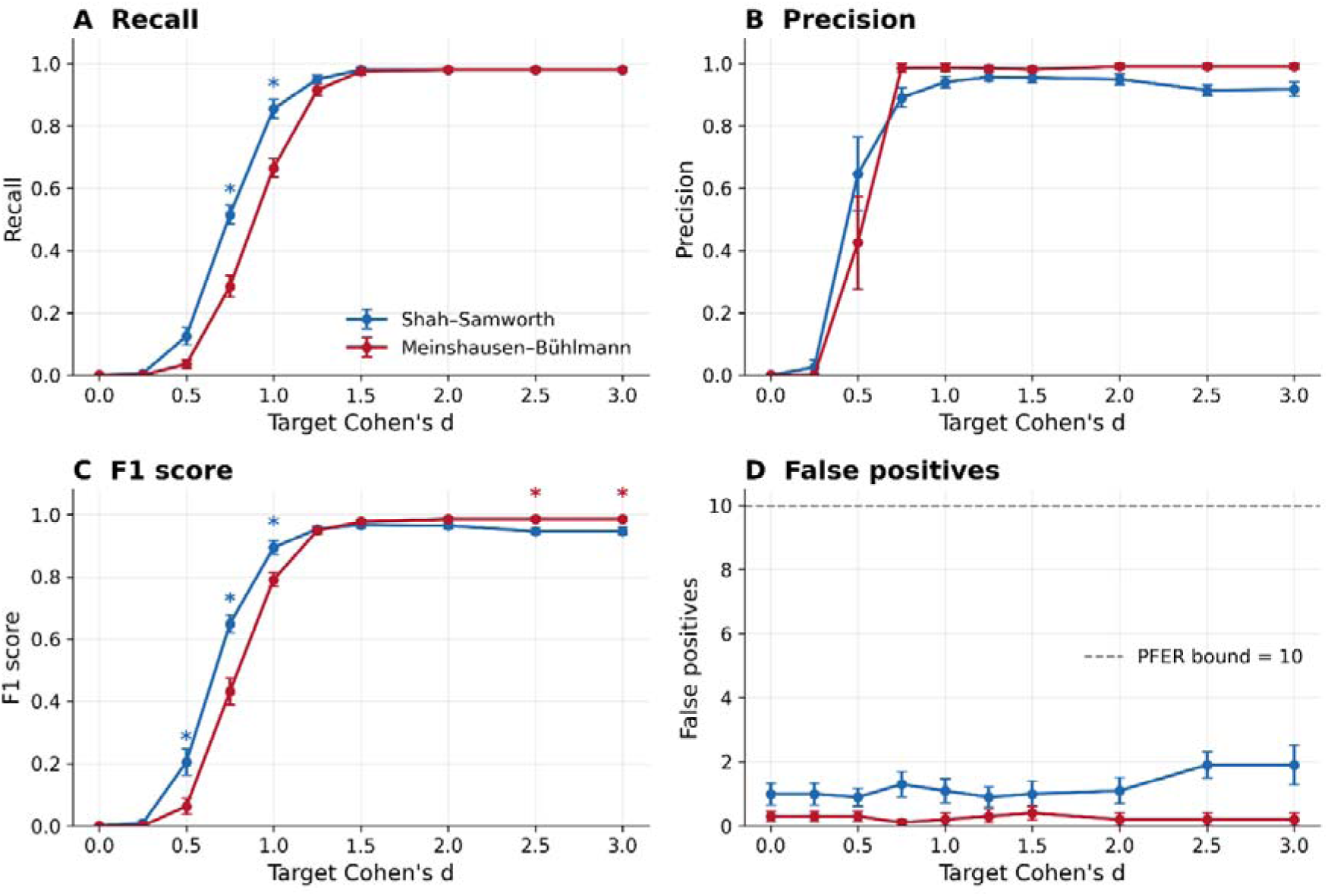
The Shah–Samworth threshold recovers more true biomarkers than Meinshausen–Bühlmann at moderate effect sizes. Recall. (A), precision (B), F1 (C) and false positives (D) versus effect size for the two thresholds at L1 = 0.5, PFER = 10, on identical synthetic datasets (mean ± SEM, 10 datasets per effect size). Shah–Samworth achieves higher recall at moderate effect sizes (d 0.5–1.0), converging with Meinshausen–Bühlmann by d ≈ 1.25; Meinshausen–Bühlmann is marginally more precise. F1 favours Shah–Samworth at moderate effect sizes and Meinshausen–Bühlmann at high effect sizes (d ≥ 2.5). Both maintain false positives well below the PFER bound. Markers: Holm-adjusted paired Wilcoxon signed-rank (* p < 0.05, ** p < 0.01), coloured by the higher-scoring threshold (blue, Shah– Samworth; red, Meinshausen–Bühlmann). Over d 0.5–2.0, Shah–Samworth recovered more biomarkers (median +0.05, p = 8×10⁻□).

### Reprocessing and filtering of pancreatic cancer discovery and validation cohorts

To display the utility of *stably* for biomarker discovery, we applied the Shah– Samworth stability selection model to a publicly available dataset comparing serum proteomes from individuals with pancreatic cancer to matched controls, which is split into discovery and validation cohorts (Byeon, et al., 2024). Raw data were reprocessed using DIA-NN (version 2.5.0). The summary statistics of both datasets are presented in Table 1. After reprocessing in DIA-NN, the discovery cohort comprised 317 protein groups across 120 samples, and the validation cohort 276 protein groups across 56 samples. Overall missingness was 18.6% (Discovery) and 11.6% (Validation), and per-protein missingness was right-skewed. Filtering for proteins with at least two proteotypic peptides retained 252 (Discovery) and 238 (Validation) protein groups; removing proteins with greater than 30% missingness left a final analysis set of 214 proteins across 120 samples (Discovery) and 206 proteins across 56 samples (Validation). Of this set, 201 protein groups were detected in both cohorts and were therefore assessable for cross-cohort stability.

**Table 1.** Summary statistics of the discovery and validation cohorts after searching in DIA-NN and filtering for high-quality proteins.

|  | **Discovery** | **Validation** |
| --- | --- | --- |
| Proteins identified | 317 | 276 |
| Samples | 120 | 56 |
| Overall missingness | 18.6% | 11.6% |
| After $\geq 2$ proteotypic peptide filtering | 252 | 238 |
| After $\leq 30\%$ missingness filtering | 214 | 206 |
| Final retained | 214 (68%) | 206 (75%) |

### Univariate statistical analysis identifies many overlapping proteins for validation

Univariate statistics are often used to identify candidate biomarkers. These methods fail to recognise relationships between proteins. We applied a *limma* analysis pipeline (version 3.64.3) to both discovery and validation cohorts to identify overlapping differentially abundant proteins. Differential abundance analysis with *limma* identified 104 proteins significant at q < 0.05 in the discovery cohort (60 PDAC versus 60 healthy controls) and 96 in the validation cohort (28 PDAC versus 28 healthy controls); 63 proteins were significant in both (Figure 4a). Among these, p value ranking was poorly matched between cohorts (Spearman r = 0.46), however effect size ranking matched well (Spearman r = 0.79), with 3/5 proteins (PIGR, SAA1, and HBA1) shared in the top 5 of both cohorts.

**Figure 4.**
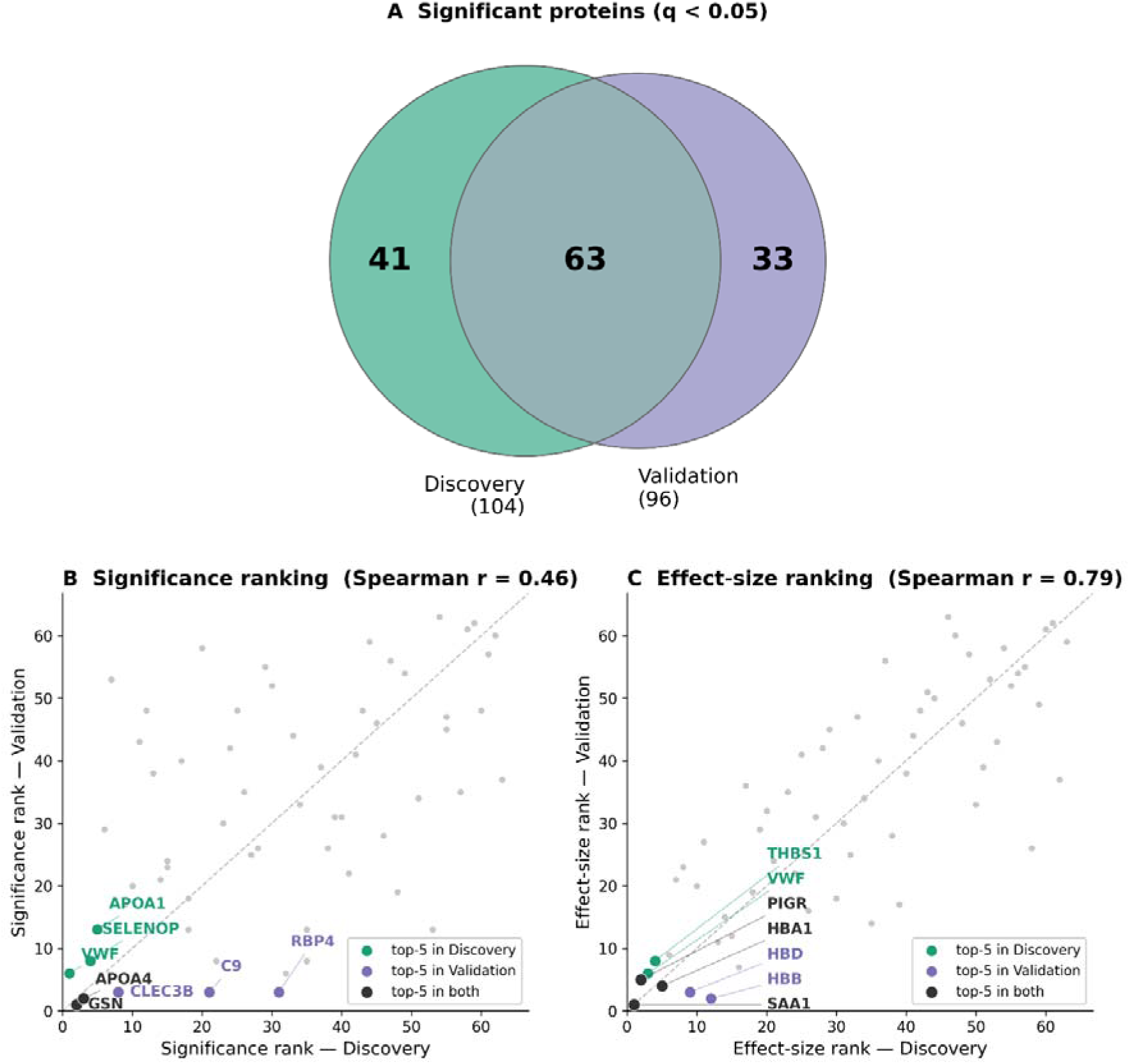
Concordance of univariate rankings between discovery and validation cohorts. (A) Proteins significant by limma (q < 0.05); 63 were significant in both datasets. Among these 63, ranking by significance (q-value) was weakly concordant across cohorts (B; Spearman r = 0.46), whereas ranking by effect size (|log□FC|) was strongly concordant (C; Spearman r = 0.79). In both panels, rank 1 is the top-ranked protein and the five top-ranked per cohort are highlighted. The difference reflects that significance ranking conflates effect size with measurement precision, so effect-size ranking more reproducibly captures the underlying signal.

### *stably* identifies stable biomarkers of pancreatic cancer in discovery and validation cohorts

We applied *stably* to the discovery and validation cohorts using an ElasticNet base learner with an L1 ratio of 0.5 and a PFER of 10, as used in the synthetic benchmarking. In the discovery cohort (60 PDAC, 60 controls), 17 proteins passed the stability threshold, with effect sizes ranging from −1.61 to +1.91 (Figure 5A). The VWF protein was the most stable feature, selected in 100% of iterations, and GSN, PIGR, F5, CFH, and C2 were each selected in >90% of iterations. In the validation cohort (28 PDAC, 28 controls), 15 proteins passed the stability threshold, of which GSN and SERPINA1 were selected in >90% of iterations, with effect sizes ranging from −1.48 to +1.17 (Figure 5B). Two proteins (GSN and APOA4) were stable in both cohorts (Figure 5C), and both shared the same directionality between cohorts (Figure 5D).

**Figure 5.**
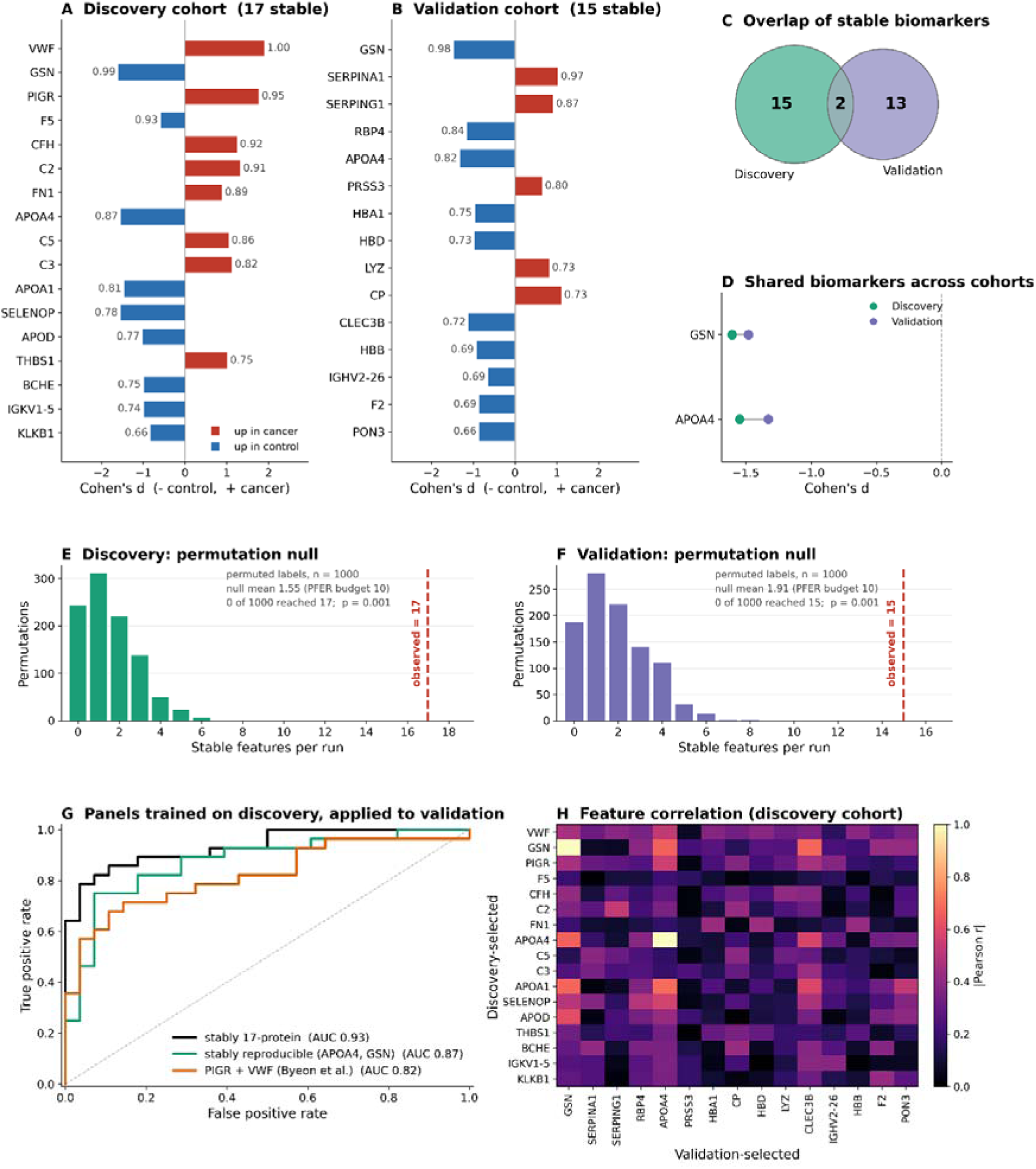
Reproducibility and predictive validation of *stably* selected biomarkers across the discovery and validation cohorts. (A, B) Stable biomarkers identified by *stably* (Shah–Samworth threshold, ElasticNet L1 = 0.5, PFER = 10) in the discovery (A; 17 features, 60 PDAC / 60 control) and validation (B; 15 features, 28 / 28) cohorts, shown as Cohen’s *d* (bars; − higher in controls, + higher in cancer), coloured by direction and ordered by selection probability. (C) Overlap of the stable sets: two proteins (GSN, APOA4) were stable in both. (D) The two shared biomarkers were concordant in direction and effect size between cohorts. (E, F) Label-permutation nulls for the discovery (E) and validation (F) cohorts. The entire selection procedure was repeated on 1000 permutations of group labels. Bars show distribution of the number of stable features per permuted run; the dashed line marks the observed value. No permutation reached the observed panel size in either cohort (17 and 15 respectively; p = 0.001). Under permutation, the procedure selected a mean of 1.55 (E) and 1.91 (F) features per run against a PFER budget of 10. (G) Panels trained on the discovery cohort, with the logistic regression regularisation strength tuned by cross-validation within that cohort and applied unchanged to the held-out validation cohort. The 17-protein discovery panel achieved AUC = 0.93, the two cross-cohort stable proteins (GSN, APOA4) 0.87, and the PIGR + VWF panel of Byeon et al., measured here in the same DIA-MS data, 0.82. The 17-protein panel significantly outperformed PIGR + VWF (paired bootstrap over the 56 validation samples, p = 0.009); no other pairwise difference reached significance. (H) Absolute Pearson correlations between discovery-selected and validation-selected proteins, computed across the 120 discovery samples. Apart from GSN and APOA4, which were stable in both cohorts and so appear on both axes at |r| = 1, no pair exceeded |r| = 0.69 and the median correlation was 0.22.

Significance was assessed by repeating the entire selection procedure on 1000 label permutations in each cohort. In the discovery cohort, no permutation produced a panel as large as the observed 17 stable features (p = 0.001, 95% CI 0.000-0.004), and a selection probability as high as the observed 1.000 in 3 out of 1000 (p = 0.004, 95% CI 0.001 – 0.009) (Figure 5E). The validation cohort behaved likewise: no permutation reached the observed 15 stable features (p = 0.001, 95% CI 0.000-0.004), and the observed maximum of 0.98 was matched or exceeded in 13 out of 1000 (p = 0.014, 95% CI 0.007 – 0.022) (Figure 5F). Under permutation, the procedure selected a mean of 1.55 (discovery) and 1.91 (validation) features per run against a PFER budget of 10. Individual stable features were common under the null: 756 of 1000 discovery and 812 out of 1000 validation permutations produced at least one, and the null median maximum selection probability (0.73 and 0.76) exceeded the respective thresholds (0.65 and 0.66).

Membership overlap between cohorts is an overly demanding measure of a biomarker’s reproducibility. A more appropriate measure is whether a panel of biomarkers maintains performance in an independent cohort. A logistic regression classifier was fitted to the 120 discovery samples using the 17 stable proteins, and imputation, scaling and regression parameters were all estimated. Applied unchanged to the validation samples, the panel discriminated patients with PDAC from healthy controls with a ROC-AUC of 0.93 (Figure 5G).

In their original analysis, Byeon et.al (2024) identified four candidate protein biomarkers in the discovery cohort, defined by each biomarker’s log_2_(FC), q value, and ROC-AUC in both cohorts. Of these four biomarkers (PIGR, fibrinogen, VWF, and CRP), all but CRP were validated by ELISA in the validation cohort, where they found that a panel of PIGR and VWF achieved a ROC-AUC of 0.90. We here use the ROC-AUC for this two-protein panel derived from the DIA-MS data for a fair comparison, where it achieved a ROC-AUC of 0.82; the comparison made here is therefore between panels measured on a common platform rather than between assay technologies. Comparing our 17-protein panel to the two-protein panel of PIGR and VWF, the 17-protein panel achieved a significantly higher ROC-AUC (0.93 versus 0.82; bootstrap difference +0.11, 95% CI +0.02 to +0.22, p = 0.009; p = 0.027 after Bonferroni correction for three comparisons). The two proteins selected as stable in both cohorts by *stably*, GSN and APOA4, achieved a ROC-AUC of 0.87, which was neither statistically greater than the panel selected by Byeon et al. (AUC 0.87 versus 0.82; bootstrap difference +0.05, 95% CI −0.08 to +0.19, p = 0.45) nor significantly worse than the full 17-protein panel (AUC 0.87 versus 0.93; bootstrap difference +0.06, 95% CI −0.02 to +0.15, p = 0.15).

ElasticNet penalisation retains only one representative from a group of correlated features, and among near-equivalent members that choice can be close to arbitrary. Limited membership overlap between cohorts could therefore be misleading: a protein stable in only one cohort might correlate strongly with one stable in the other. Here, however, correlations between the discovery-selected and validation-selected proteins were poor (Figure 5H). Apart from GSN and APOA4, which were stable in both cohorts, no pair exceeded |r| = 0.69, and the median correlation across the matrix was 0.22. This is against a background in which 60.7% of proteins correlate at |r| > 0.5 with at least one other. The proteins selected in only one cohort are therefore not interchangeable proxies for those selected in the other, indicating that the limited membership overlap reflects genuine disagreement between the cohorts rather than arbitrary substitution within correlated clusters.

## Discussion

Machine learning for the identification of biomarkers in untargeted proteomics data allows deeper probing to identify biomarkers of disease for validation. This growing dimensionality has motivated adoption of a range of feature selection methods and aims to isolate a discriminatory subset of proteins. Wrapper approaches such as Boruta use a multiply trained random forest to identify features that are significantly better predictors of group membership than randomised shadow features (Kursa and Rudnicki, 2010), while recursive feature elimination iteratively discards the least informative features from a base learner such as a support vector machine (Guyon, et al., 2002). Embedded methods, including the feature importances of random forests and gradient-boosted trees, or the non-zero coefficients of penalised regression are also widely used to rank and select features. While these methods are effective at producing a predictive subset, they share two limitations in the context of biomarker discovery: the selected set is sensitive to data splitting and resampling, and, critically, none place a formal bound on the number of expected false positives within the returned panel. A method may therefore identify a highly predictive set of proteins yet does not offer a guarantee on how many of those proteins are redundant, an unsuitable foundation for committing resources to downstream validation.

Among multivariate feature-selection approaches, two prominent frameworks attach a formal error guarantee to the selected feature set. The knockoff framework constructs synthetic variables that reproduce the dependence structure of the measured features and controls the false discovery rate by comparing the importance of each real feature with its knockoff counterpart (Candès, et al., 2018; Rina Foygel and Emmanuel, 2015). However, the fixed-design formulation imposes sample-size and rank requirements that are rarely met in untargeted proteomics, while the model-X extension requires the joint distribution of the proteins to be known or accurately estimated, which is fragile in the small-sample, densely correlated setting characteristic of proteomic data (Figure 2). Moreover, false discovery rate control bounds the expected proportion of false positives among selected features, a guarantee that is less directly interpretable for a small, fixed biomarker panel than a bound on the expected number of false positives.

Stability selection provides this latter guarantee. By repeatedly subsampling the data and retaining only those features selected with high frequency, it bounds the PFER while requiring weaker assumptions than knockoffs (Meinshausen and Bühlmann, 2010; Shah and Samworth, 2013). This makes it well suited to small, correlated proteomics data, as demonstrated by the false-positive counts maintained well below the PFER bound across all effect sizes in our synthetic benchmark (Figure 3D), and confirmed on real, densely correlated proteomic data by label permutation (Figure 5E, F). Despite this, existing implementations such as the *stabs* and *sharp* R packages, or the now-deprecated scikit-learn stability-selection module, require considerable coding expertise and are not tailored to the output of DIA proteomics workflows. *stably* addresses this gap by bringing error-controlled stability selection directly to DIA-NN-processed data, representing the first implementation of a proteomics-specific stability selection package for the processing of untargeted proteomics data, requiring minimal coding experience to use.

Both the Meinshausen–Bühlmann and Shah–Samworth models bound the number of false positives, but they differ in how they convert a target error rate into a selection-probability threshold. Meinshausen–Bühlmann holds only under the assumptions that all noise proteins share an equal selection probability and that signal proteins are selected more frequently than noise, conditions unlikely to be met in proteomic data where the dense correlation between proteins means noise proteins correlated with informative features are not selected with equal probability (Figure 2C). The Shah–Samworth model removes these assumptions (Shah and Samworth, 2013). It instead constrains the shape of the distribution of selection proportions: the function D(η, t, N, r) computes the worst-case probability that a noise protein’s selection frequency exceeds the threshold, given its expected selection probability and an assumption of r-concavity (with r = −1/2 and r = −1/4 entering the bound). By limiting how much probability mass can concentrate near the threshold, these shape constraints yield a less conservative bound, and therefore a lower selection-probability threshold for the same false-positive guarantee. This is confirmed in our synthetic benchmarking, where at a PFER target of 10, Shah– Samworth set a threshold of 0.54 against 0.71 for Meinshausen–Bühlmann (Figure 3).

This lower threshold is the source of Shah–Samworth’s advantage; weaker-signal biomarkers are recovered, and this gain is concentrated at the moderate effect sizes (d ≈ 0.5–1.0) typical of biomarker discovery; the two frameworks converge once the signal clears both thresholds (Figure 3A, C). Notably, the more conservative Meinshausen–Bühlmann threshold did not translate into a failure of error control on data. It produced fewer false positives than Shah–Samworth (0.3 versus 1.0), with both below the PFER target of 10 (Figure 3D). The assumption-free Shah–Samworth guarantee is therefore preferable not because the Meinshausen–Bühlmann bound breaks down in practice, but because it delivers the same formal control under weaker requirements while recovering more true biomarkers. For this reason, we adopted the Shah–Samworth threshold as the default in *stably*.

Univariate differential abundance analysis is the most common route to candidate biomarkers for validation. Applying this to a publicly available dataset, *limma* returned 63 proteins significantly differentially abundant in both the discovery and validation cohorts (Figure 4A). A limitation of this approach is the absence of a principled way to convert a long, ranked list into a tractable panel for downstream testing. A significance threshold identifies which proteins differ between groups but offers no indicator of how well that protein predicts group membership, no error-controlled cutoff on how many proteins to carry forward, no protection against the redundancy from co-regulated proteins, and a ranking that is itself unstable between cohorts. Ranking by significance was only weakly reproducible across cohorts (Spearman r = 0.46), although ranking by effect size was better (r = 0.79; Figure 4b, c). A practical consequence is that, where univariate ranking is used to prioritise candidates for downstream validation, effect size is a more reproducible criterion than significance; but neither resolves the more fundamental problems of panel size and false-positive control.

The 17-protein panel generated on the discovery cohort achieved significantly greater performance than the panel selected by Byeon et.al (AUC 0.93 versus 0.82, Figure 5G, p = 0.009. Permutation testing by random label shuffling confirmed the significance of both panels: no permutation of either cohort produced a panel as large as the one observed (Figure 5E, F). This complements the previous synthetic benchmarking of *stably* by showing the PFER bound holding in complex proteomics data, where permuted runs selected a mean of 1.55 (discovery, Figure 5E) and 1.91 (validation, Figure 5F) features — well below the PFER budget of 10. Substantial numbers of features were nonetheless selected as stable individually under the null, and the null median maximum selection probability sat comfortably above the selection threshold in both cohorts (0.73 against a threshold of 0.65 in discovery; 0.76 against 0.66 in validation). This highlights the importance of considering biomarkers as a panel, where panel size, rather than the stability of any single protein, carries the weight of the discovery.

Notably, the two candidate biomarker proteins for validation identified across cohorts by *stably* was supported by the univariate analysis, but not in a way that makes them obvious candidates. GSN and APOA4 were among the 63 proteins significantly differentially abundant in both cohorts. However, they were embedded within a much larger jointly significant set, and any decision to prioritise them for validation from the univariate results would have required an additional, arbitrary filtering or ranking rule. By contrast, *stably* selected GSN and APOA4 directly as the only proteins stable in both cohorts (Figure 5C), and these two proteins as a panel discriminated pancreatic cancer from control in the validation cohort with performance statistically indistinguishable from a full ML-derived 17-protein discovery panel (AUC 0.87 versus 0.93; Figure 5G, p = 0.15). Univariate testing answers which proteins differ between groups, whereas stability selection identifies a compact, reproducible, error-controlled panel suitable for downstream validation.

It is notable that only two proteins were also stably selected across cohorts, out of panels of 17 and 15 proteins respectively (Figure 5C). Whether reproducible stable selection across cohorts is itself important is debatable. The operative endpoint for a biomarker panel is not repeated selection in a second global -omics cohort, but predictive performance in an independent cohort i.e. whether, when applied to new patients, it achieves sufficient sensitivity and specificity in identification of patients within a target group. On this measure, the discovery panel transferred well to a second set of patient samples (validation AUC 0.93; Figure 5G). This panel achieved significantly greater ROC AUC than the panel selected by Byeon et.al (AUC 0.93 versus 0.82, p = 0.009, Figure 5G), displaying the utility of *stably* as a superior biomarker discovery tool to conventional univariate analyses in DIA-MS data. A two-protein panel of the reproducibly selected core (APOA4, GSN) performed comparably (AUC 0.87; ΔAUC 0.06, p = 0.15), suggesting that in these cohorts the remaining 15 proteins add no substantial additional discrimination, though the modest validation cohort size (n = 56) leaves open the question as to whether they contribute at higher power. The reproducible finding is therefore the core differentiating markers that translate. *stably* accordingly functions as a discovery tool that identifies stably selected signals from which candidate multi-biomarker panels can be built for downstream verification in a targeted assay. Its output represents targets whose selection is robust across data subsamples rather than contingent on a single partition, with explicit control of the expected number of false positives – providing a stronger starting point for clinical translation.

## Limitations

The synthetic benchmark, while allowing false-positive control to be assessed against a known ground truth, does not fully reproduce the structure of real proteomic data. In particular, the synthetic proteins were far less correlated than those in the discovery cohort (1.3% versus 60.7% of proteins strongly correlated with at least one other; Figure 2c) and the data were correspondingly higher-dimensional. The benefit of an ElasticNet base learner in proteomics data is for discerning between correlated biomarkers, and in the context of stability selection, choosing the same biomarker repeatedly from that group. The benchmark using synthetic data with limited correlation structure provides a conservative test of feature recovery but is an imperfect proxy for how the method behaves on densely correlated proteomes. Despite this, the permutation testing in Figure 5E and F shows that false positives remain well below the PFER bound on real, densely correlated proteomic data. The generative model also imposes effect sizes as simple shifts on log-normal intensities with an intensity-dependent missingness mechanism, and so does not capture the batch effects, non-linear relationships, or pathway-level co-regulation present in biological data. Statistical comparison of the two thresholds was focused on a proteomics-typical range of effect sizes (d = 0.5–2.0), although the benchmark spanned d = 0–3.0; recall was poor at smaller effect sizes, reflecting a requirement for adequate effect size and sample size that stability selection does not remove. A limitation with the validation cohort is the size, at only 56 samples, the comparisons between panels carry wide confidence intervals and are not powered to establish equivalence. The non-significant differences reported can therefore be read as an inability to distinguish the panels, or as evidence that they perform identically.

## Acknowledgements

This work was funded by UK Research and Innovation through a Medical Research Council Doctoral Training Programme award (MR/W007428/1). The work was facilitated by the Manchester National Institute for Health Research Biomedical Research Centre and the Greater Manchester Comprehensive Local Research Network. The authors acknowledge the use of Claude code to aid code development.

## Code availability

*stably* is available at (https://github.com/byrnedaniel5-eng/stably), along with detailed guides on how to download and analyse proteomics datasets. The version of *stably* used in this paper is archived at PyPI (https://pypi.org/project/stably/).

## Data availability

Data are available at the ProteomeXchange (PXD046438) for the pancreatic cancer discovery and validation cohorts.

## Conflict of interest

The authors declare no conflict of interest.

## Notes

https://proteomecentral.proteomexchange.org/cgi/GetDataset?ID=PXD046438-1&test=no

### Competing Interest Statement

The authors have declared no competing interest.

